# Chronic stress triggers seeking of a starvation-like state in anxiety-prone female mice

**DOI:** 10.1101/2023.05.16.541013

**Authors:** Hakan Kucukdereli, Oren Amsalem, Trent Pottala, Michelle Lim, Leilani Potgieter, Amanda Hasbrouck, Andrew Lutas, Mark L. Andermann

## Abstract

Elevated anxiety often precedes anorexia nervosa and persists after weight restoration. Patients with anorexia nervosa often describe hunger as pleasant, potentially because food restriction can be anxiolytic. Here, we tested whether chronic stress can cause animals to prefer a starvation-like state. We developed a virtual reality place preference paradigm in which head-fixed mice can voluntarily seek a starvation-like state induced by optogenetic stimulation of hypothalamic agouti-related peptide (AgRP) neurons. Prior to stress induction, male but not female mice showed mild aversion to AgRP stimulation. Strikingly, following chronic stress, a subset of females developed a strong preference for AgRP stimulation that was predicted by high baseline anxiety. Such stress-induced changes in preference were reflected in changes in facial expressions during AgRP stimulation. Our study suggests that stress may cause females predisposed to anxiety to seek a starvation state, and provides a powerful experimental framework for investigating the underlying neural mechanisms.

## Introduction

Anorexia nervosa has the highest mortality rate of any psychiatric disorder.^1, 2^ Key physical signs and symptoms include willful self-starvation through a combination of dieting, fasting, and excessive exercising.^3–5^ Notably, the prevalence of this disorder is five to ten times higher in females than in males.^6^ Moreover, depression, anxiety, and obsessive-compulsive disorder (OCD) are common among anorexia nervosa patients. Despite the severity of this disorder, current treatment options, such as psychotherapy and nutritional rehabilitation, are expensive and often ineffective.^2, 3, 6–8^

While anorexia nervosa has been documented for over 300 years,^9^ the underlying causes remain unknown. In particular, it is unclear why individuals with anorexia nervosa continuously engage in behaviors that maintain a long-term starvation state despite being severely underweight.^3, 10, 11^ Current behavioral and genetic models in mice have advanced our understanding of certain physiological and behavioral features of anorexia nervosa^12^, but they fail to capture a key hallmark of this disorder – willful starvation.^4, 12^ For example, the widely-used activity-based anorexia (ABA) model combines calorie restriction and scheduled feeding with free access to a running wheel to elicit anorexia nervosa phenotypes such as chronic weight loss and hyperactivity.^13, 14^ A second mouse model^15^ involves early-life stress and a genetic mutation that is associated with elevated anxiety and increased severity of anorexia nervosa.^15–18^ Notably, these mice exhibit brief, sporadic bouts of willful food restriction. However, the model suffers from a high mortality rate and does not capture the persistent performance of deliberate actions that maintain a starvation state. Thus, there is a pressing need for a pre-clinical mouse model that captures the intentional seeking of a starvation state and the dependence of this behavior on baseline anxiety levels.^19–21^

In healthy individuals, hunger is considered a mildly aversive state that drives food consumption to maintain energy homeostasis.^22, 23^ In mice, artificial activation of neurons in the arcuate nucleus of the hypothalamus that express agouti-related peptide (AgRP) and neuropeptide Y (NPY) is sufficient to drive seeking and consumption of food.^24–27^ Additionally, male mice voluntarily avoid the artificial starvation-like state induced by stimulation of AgRP neurons, suggesting that it is aversive.^26^ Notably, food restriction and stimulation of AgRP neurons can both reduce anxiety-like behaviors and sensitivity to subsequent acute stressors, thereby promoting active exploration and reducing neural sensitivity to stressful cues and threatening environments.^28–35^

This ability of food restriction and the associated elevation in AgRP neuron activity to suppress anxiety in certain contexts may explain why some individuals with anorexia nervosa find hunger to be pleasant.^4, 36, 37^ Indeed, the on-set of this disorder is often preceded by childhood trauma, stressful early life experience, or social stress.^4, 8, 38^ We hypothesized that, in contrast to healthy individuals, some individuals exposed to high levels of anxiety and stress would begin to willfully seek a state of starvation as a means of coping with stress and reducing anxiety. Below, we tested this hypothesis in transgenic mice by modeling the willful seeking of a starvation state using a paradigm for voluntary self-stimulation of AgRP neurons.

## Results

### A new behavioral model for voluntary seeking of a starvation-like state in mice

We developed a behavioral paradigm to test whether mice intentionally seek a starvation-like state following repeated exposure to a stressful environment. To this end, we designed a virtual reality (VR) place preference protocol in head-fixed mice to improve behavioral and stimulus control. In contrast to traditional place preference assays in freely moving mice, this VR place preference assay enables concurrent tracking of facial expressions (see Figure 4) and, in the future, will facilitate concurrent two-photon calcium imaging, high-density electrophysiology, and recordings of autonomic signals. Our protocol consisted of multiple experimental phases (Figure 1A; >1,000 hour-long sessions, 37 sessions/mouse across 15 male and 17 female mice). This allowed us to gain a better understanding of the variability in behavior within and across mice.

**Figure 1.**
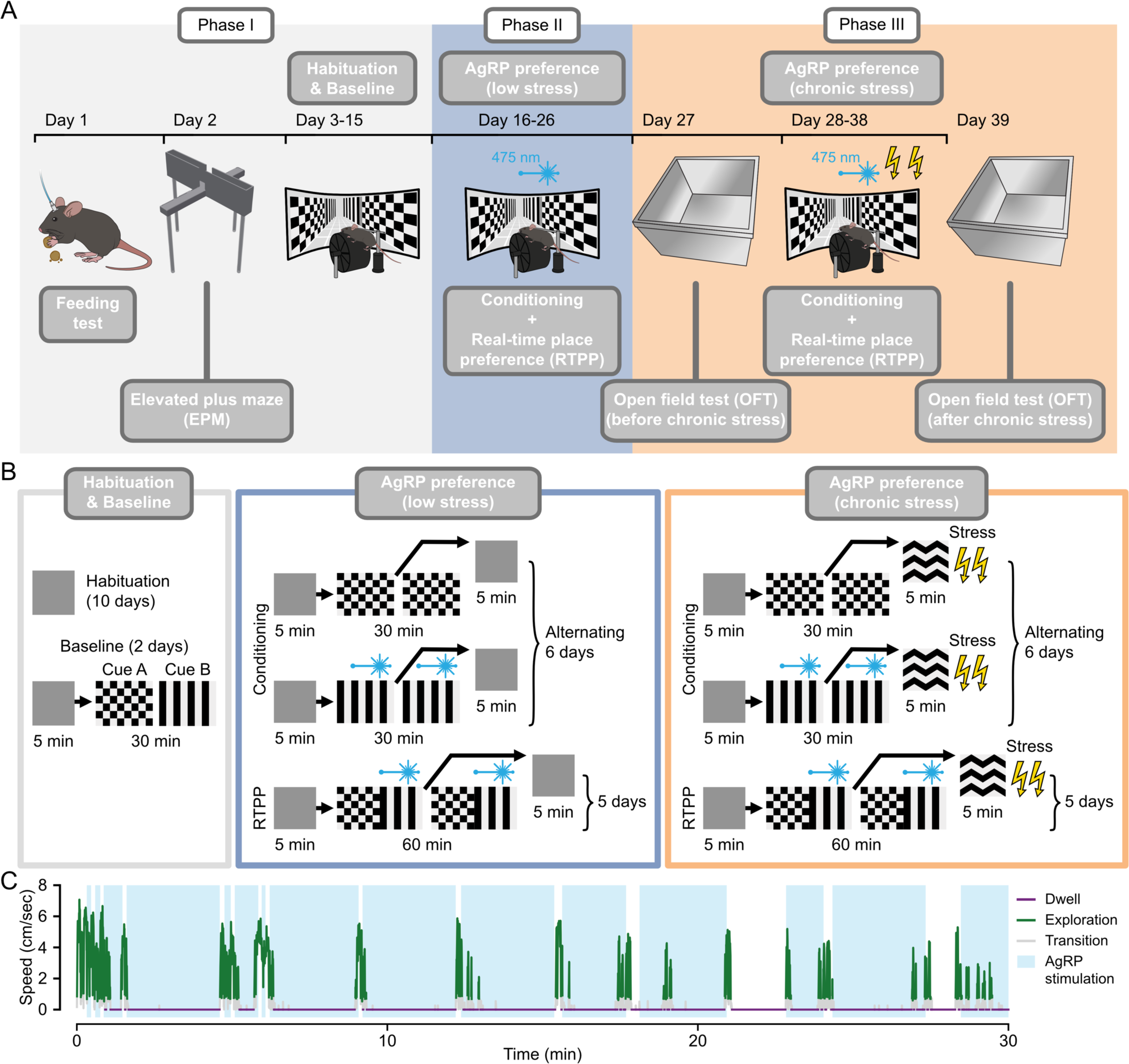
A head-fixed behavioral paradigm in virtual reality to study voluntary seeking of a starvation-like state in mice. **(A)** The experimental timeline consisted of three phases. *Phase I* (gray box) began with confirming the efficacy of AgRP neuron stimulation by measuring home-cage food intake in response to optogenetic stimulation. Then, baseline anxiety-like behavior was assessed in an elevated plus maze (EPM) assay. Finally, mice were extensively habituated to head fixation and to the virtual reality (VR) environment prior to assessing their initial bias toward visual cues (see below). In *Phase II* (blue box), following conditioning sessions in which AgRP stimulation was paired with one visual cue and no stimulation was paired with the other cue, mice were tested for their preference for AgRP stimulation in a real-time place preference (RTPP) assay. In *Phase III* (orange box), *Phase II* was repeated during chronic stress induced by a five-minute period of mild tail shocks within every session, and verified by comparing open field tests (OFT) conducted before and after the *Phase III* sessions. **(B)** *Gray box, top*: mice were habituated in a corridor with gray walls (to avoid desensitization to visual cues) for 10 days before pairing with stimulation. *Gray box, bottom*: two distinct, luminance-matched visual cues were presented on the walls of the corridors: checkers (cue A) and stripes (cue B). Following habituation, the initial bias for the cues was assessed over two days. The least preferred cue was paired with AgRP stimulation for the rest of the experiment (Methods). *Blue box, top*: mice were exposed for 30 minutes to either a cue paired with AgRP stimulation (stim), or to a second cue not associated with stimulation (neutral) on alternating sessions (6 sessions total). *Blue box,* bottom: this was followed by the RTPP assay (five onehour daily sessions) to assess preference for AgRP stimulation under low-stress conditions. *Orange box,* bottom: following another 6 days of conditioning (three days per cue) involving chronic stress (see below), mice were tested for their preference for AgRP stimulation for 5 days. *Orange box, top*: chronic stress was induced by five minutes of repeated mild tail shocks in a corridor with a distinct third visual cue (orange box, right). **(C)** Example trace of locomotion speed, illustrating how an example mouse (female, post-stress; AgRP preference index: 0.83) navigates in the VR environment. Behavior was categorized into three states: *dwell* (purple), when mice were stationary; *explore* (green), when mice were actively exploring the virtual environment; and *transition* (gray), an intermediate speed at transitions between the other states. The example mouse actively explored both cues but chose to spend more time dwelling in the corridor paired with AgRP stimulation. See also Figure S1.

We used optogenetic stimulation of AgRP neurons in the arcuate nucleus of the hypothalamus to induce a starvation-like state, as AgRP neuron activity is elevated during starvation states.^26, 39^ Further, AgRP activation drives a coordinated set of behaviors associated with starvation, including voracious feeding and a reduction in sensitivity to pain, acute stressors and anxiety.^24, 25, 29– 32, 40, 41^ To reduce variability in stimulation strength across mice, we used a transgenic approach involving uniform expression of channelrhodopsin (ChR2) in all AgRP neurons in *Agrp*-IRES-Cre::ChR2-EYFP mice.^42, 43^ Photostimulation was achieved via an optic fiber placed above the arcuate nucleus (Figures S1A-S1B). Effective stimulation of AgRP neurons was confirmed by comparing home-cage food intake during one hour of AgRP stimulation in mice with *ad libitum* access to food during the light cycle (Figure S1C), when mice normally eat very little. During *Phase I* of the protocol, we assessed the baseline level of anxiety-like behavior using an elevated plus maze (EPM; see Figure 3 below). To reduce any stress related to head fixation and exposure to the VR setup, we gradually habituated mice to increasing levels of head fixation for ten days prior to the experiment, which effectively mitigates restraint stress.^44–46^ *Phase I* concluded with the assessment of any weak baseline preferences towards one of two virtual corridors distinguished by their visual cues (Figures 1A and 1B; gray box).

In *Phase II*, these cues were then paired with either the presence (stimulation) or absence (neutral) of AgRP stimulation (Figure 1B; blue and orange boxes). The slightly less preferred cue was paired with AgRP stimulation. This same pairing was maintained throughout the experiment. Because AgRP stimulation takes several tens of seconds to minutes to drive food-seeking behavior^24, 25^, we carried out six daily baseline conditioning sessions, consistent with a prior study.^26^ On alternating days, mice were exposed to the corridor containing the cue paired with AgRP stimulation or the neutral cue (Figure 1B; blue box). We then assessed realtime place preference (RTPP) during five additional daily sessions in which mice could choose to dwell in a corridor containing the stimulation cue paired with AgRP stimulation or the neutral cue (Figure 1A; *Phase*). Mice had *ad libitum* access to food except during and in the 30 minutes following each session to avoid prolonged effects of AgRP stimulation on food intake.

To test the hypothesis that mice may seek activation of AgRP neurons following chronic stress, we repeated the sequence of six conditioning sessions and five RTPP sessions, but this time with exposure to repeated tail shocks during a 5-minute period halfway through each session (Figure 1A; *Phase III*). During this period, a distinct visual cue was presented (Figure 1B, orange box) while the mouse received mild tail-shocks with unpredictable timing (2-3 s train of 12-17 0.35 mA pulses every 30-35 s; pulse duration: 50 ms). The efficacy of chronic stress induction was assessed using open-field tests (OFT) performed before and after the series of sessions containing shocks (Figure 1A; *Phase III*).

Mice actively navigated the virtual reality environment and often spent more time in a stationary state (i.e. dwelling) in locations associated with their preferred cue in the virtual corridor. This is evident in the example mouse in Figure 1C that chose to dwell in the corridor associated with AgRP stimulation. Because such dwell periods could be long-lasting, thereby reducing the number of active choices to stay in either corridor, we leveraged the VR setup to ‘teleport’ the mouse to the opposite virtual corridor if it remained in one corridor for over three minutes. Within seconds of forced teleportation to the neutral corridor (and associated cessation of AgRP stimulation), this same mouse began to run through this new corridor until it returned to the AgRP stimulation corridor (Figure 1C). For all subsequent analyses, we excluded bouts of locomotion (related to exploration and transition between corridors) and calculated a preference index for each mouse based on the relative duration that it dwelled in a stationary state in the AgRP stimulation corridor versus in the stationary corridor, ranging from 1 (always) to -1 (never).

### AgRP stimulation is aversive in male

According to the drive reduction theory, hunger is an aversive state, and animals eat in order to alleviate this state.^26, 27^ In line with this, activation of AgRP neurons was previously shown to be mildly aversive in male mice.^26^ However, other studies also suggest that AgRP stimulation can be anxiolytic.^28–30, 40^ Thus, we hypothesized that a strongly aversive and anxiety-promoting state such as chronic stress might, in some cases, reinforce the seeking of this starvation-like state.

We first estimated the preference of each mouse for AgRP stimulation in the absence of stress (Figures 1A-1B; *Phase I*). After conditioning the mice separately with the paired cue or unpaired cue on alternating days (Figures 1A-1B; *Phase II*), we assessed the preference for AgRP stimulation by quantifying the fraction of time spent dwelling in locations associated with each cue (see Figure 1C for an example). Consistent with previous results,^26^ male mice tended to avoid the corridor paired with AgRP stimulation, suggesting that this starvation-like state is mildly aversive (Figures 2A, 2C, and 2E). This aversion was significant on many individual sessions (Figure S2A-S2D; comparison of preference index to shuffled preference distributions, see Methods), and it was consistent across days for most male mice (Figures 2C and S2A-S2D).

**Figure 2.**
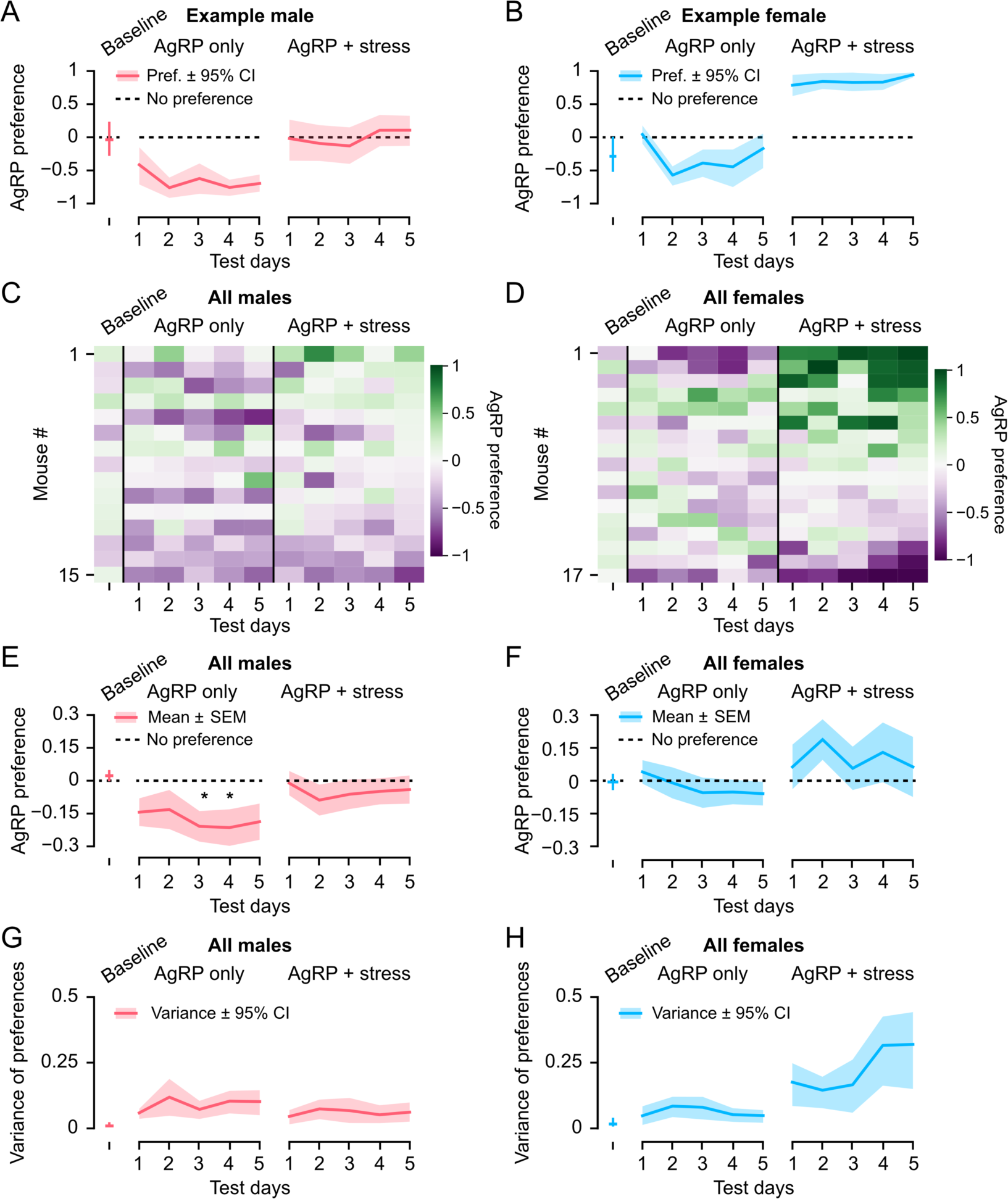
Sex-dependent polarization of preferences following chronic stress. **(A-B)** Preference of two example mice across days. **(A)** An example male mouse showed aversion to AgRP stimulation before stress (*AgRP only*). **(B)** An example female mouse showed no aversion to AgRP stimulation before stress (*AgRP only*). This mouse began to strongly prefer AgRP stimulation following conditioning sessions involving induction of chronic stress (*AgRP* + *stress*). Data represent mean ± 95% CI. **(C-D)** Heatmaps showing preferences for AgRP stimulation for all male mice (C; n=15) and female mice (D; n=17) across days. Heatmap rows are sorted by AgRP preference on Day 5 after stress (*AgRP* + *stress*). **(E-F)** Mean preferences for AgRP stimulation of male (E) and female (F) mice (n = 15 males, n = 17 females). * *P* < 0.05, one-tailed t-test with Bonferroni correction. Data represent mean ± SEM. **(E-F)** Plots of variance in preference across male mice (G) and female mice (H). Data represent mean ± 95% CI. See also **Figure S2**.

In contrast to males, females did not, on average, show a uniform aversion to or preference for AgRP stimulation prior to chronic stress (Figures 2B, 2D, 2F, S2B and 2SD). Further, while a minority of females did exhibit strong aversion in individual sessions (Figures 2D, S2B and S2D), this was less common than in males (Figures 2C, S2A and S2C).

Further analyses confirmed this sex difference in the aversion to AgRP stimulation in the absence of stress. As a measure of the efficacy of stimulation, we calculated the quantity of food consumed during one hour of AgRP stimulation in the home cage prior to any experimental sessions.^24, 47^ We observed a positive correlation between optogenetically evoked food consumption in the home cage and aversion to AgRP stimulation in subsequent VR sessions in males (Figure S2E). By contrast, this relationship was absent in females, despite a similar range and variability in preferences for AgRP stimulation (Figures 2SF and 2G-H). Moreover, both males and females consumed similar amounts of food during AgRP stimulation at the beginning of *Phase I* and after the completion of *Phase III*, suggesting similar stimulation efficacy in males and females (Figure S2G). Overall, these results suggest that the motivation to consume food in order to reduce the aversive aspects of elevated AgRP neuron stimulation may be more prominent in male mice than in female mice.

### Chronic stress polarizes preferences for AgRP stimulation in females

Next, we tested the hypothesis that repeated stressful experiences will cause some mice to voluntarily seek the starvation-like state associated with AgRP neuron stimulation (Figures 1A; Phase III). To verify that we effectively induced chronic stress, we assessed anxiety-like behavior using OFT^48, 49^ on separate days before and after this second series of conditioning and RTPP sessions, each containing a 5-minute period of unpredictable tail shocks halfway through the session. Indeed, mice consistently spent less time in the center zone of the test chamber after 11 sessions involving exposure to repeated tail shocks (Figures S2G-S2H), confirming effective induction of chronic stress.

Aversion to AgRP stimulation was, on average, blunted in male mice after induction of chronic stress (Figures 2C, 2E and S2C). Strikingly, female mice on average *preferred* AgRP stimulation after induction of chronic stress (Figures 2D, 2F and S2D). In particular, a strong preference for AgRP stimulation emerged among several females after but not prior to chronic stress (Figures 2B, 2D, and S2B; see Figure 1C for an example). This preference was consistent across days in these females, even though some of the same mice demonstrated mild aversion to AgRP stimulation prior to chronic stress (Figures 2D and S2B). These findings support our hypothesis that some individuals, particularly females, may begin to prefer a starvation state when faced with chronic stress, consistent with the higher prevalence of anorexia nervosa in the female population.

Curiously, a smaller subset of female mice showed enhanced aversion to AgRP stimulation (Figures 2D and S2B and S2D). Together, these findings indicate a polarization of preferences across females following chronic stress (Figure 2D). This polarization was evident in the marked increase in the variance of preferences for AgRP stimulation across females (Figure 2H), but not across males (Figure 2G).

### Predisposition to stress predicts individual preferences for AgRP stimulation in females

Given our finding that stress leads to increased inter-individual variability in preferences for AgRP stimulation in female mice, we explored whether any traits might predispose a subset of female mice to prefer AgRP stimulation following chronic stress, akin to previous examinations of traits that predispose humans to anorexia nervosa^50^. Although resilience to stress is associated with reduced symptoms in patients with anorexia nervosa,^51^ the retrospective nature of clinical studies prevents definitive assessment of whether a predisposition to stress-induced anxiety precedes disease onset. In contrast, our behavioral task allowed us to test for potential links to a mouse’s predisposition to stress prior to all experimental sessions. Specifically, we used the EPM to assay anxiety-like behavior and stress resilience^52, 53^ before exposing the mice to head fixation, to the VR environment, or to chronic tail-shock stress. This assay quantifies the ratio of time spent in the open arm versus the closed arm of the maze (Figures 3A and S3A. We observed greater variance in open-arm occupancy among females than males, suggesting that female mice exhibit greater inter-individual variability in their predisposition to stress (Figures 3A-3B).

**Figure 3.**
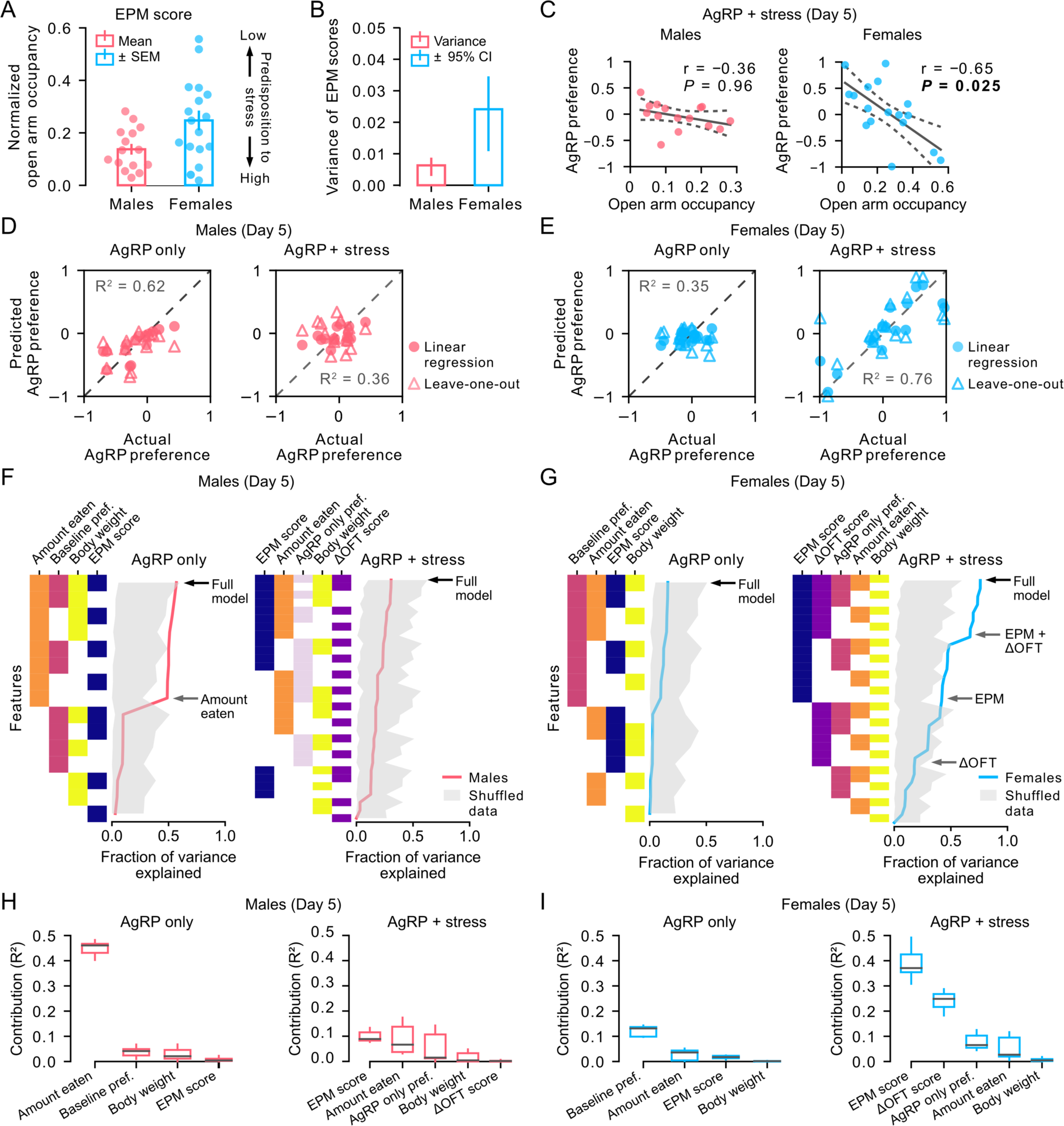
Variability in AgRP preference is predicted by predisposition to stress in females. **(A)** Open arm occupancy scores of male (left) and female (right) mice during the elevated plus maze (EPM) assay. Higher open arm occupancy is associated with a lower predisposition to stress. Data represent mean ± SEM. **(B)** Variance of open arm occupancy scores in EPM. Data represent mean ± 95% CI. **(C)** The relationship between preference for AgRP stimulation and the open arm occupancy score in males (left) and females (right) (n = 15 males, n = 17 females). r denotes Pearson’s correlation coefficient; dashed lines represent the 95% confidence band for the linear fit. *P* < 0.05 is shown in bold; *P* values are corrected for multiple comparisons using Bonferroni correction. Each circle represents a single mouse. **(D-E)** Scatterplots of predicted and actual preference for AgRP stimulation on Day 5 from a multiple linear regression model. Separate models for preferences prior to (*AgRP only*) or following (*AgRP* + *stress*) chronic stress were constructed using 4 or 5 variables, respectively. Leave-one-out analysis was performed in order to cross-validate our linear model (separate models were constructed after leaving a single mouse out of the model; see Methods). Predictions from leave-one-out analysis (open triangles) can be compared to the predictions from the multiple linear regression model (filled circles). R2 values indicate the fraction of explained variance in multiple regression models. Each data point represents a single male (D) or female (E) (n = 15 males, n = 17 females). **(F-G)** Models constructed with subsets of variables used in the full model (top row; black arrows). Fraction of explained variance (R2, solid line) is plotted for males (F) and females (G) as compared to the 95th percentile of the shuffle controls (gray shading). Models that contain one or two variables that contribute substantially to the explained variance of the full model are denoted with gray arrows. Variables are sorted by the amount of variance explained (see below). **(H-I)** Box plots of the contribution of individual variables to the variance explained (R2) by the full model for male and female mice (n = 15 males, n = 17 females). The contribution of each variable was calculated by subtracting the R2 value of the model without that variable from the full model. Variables are sorted by the amount of variance explained. Box plots represent median, 2nd and 3rd quartiles, whiskers represent minimum and maximum. See also **Figure S3**.

We examined whether this predisposition to stress might predict the preferences of individual females for AgRP stimulation after chronic stress. We focused on the final day of testing (Day 5) when cumulative prior exposure to stressors was greatest. Strikingly, we found a strong relationship in females between preference for AgRP stimulation and EPM scores estimated weeks earlier, prior to chronic stress (Figure 3C; right, and Figure 1A; experimental timeline). Specifically, females that preferred AgRP stimulation after chronic stress spent less time in the open arm of the EPM, indicating that they were relatively more anxious or less tolerant to stress even before the start of the experiment. This strong correlation was evident throughout the five days of testing following chronic stress (Figure 3C; right, and Figure S3C). In contrast, no such relationship between EPM behavior and AgRP preference was observed in males (Figure 3C; left, and Figure S3B). The absence of such correlation in males prior to or after chronic stress, or in females prior to chronic stress, is unlikely to be due to lower variance in AgRP preferences. Indeed, we observed a similarly strong relationship in males prior to stress between AgRP preference and food consumed in the home cage during AgRP stimulation (Figure S2D).

We considered whether AgRP preference in females might depend on phase within the estrous cycle. Differences in circulating estrogen across the estrous cycle can affect the drive to consume food.^54, 55^ While AgRP neurons can drive food intake across each phase of the estrous cycle, the potency of AgRP stimulation may depend on the estrous phase.^56^ While we did not track the estrous cycle in our female mice, we considered its potential impact as follows: given that the mouse estrous cycle lasts 4-5 days,^57, 58^ we reasoned that the mice should be in estrous at least on one day within our protocol of 5 consecutive days of estimation of RTPP. Yet, the sign of AgRP preference was consistent across days following chronic stress (Figures 2C, S2A, and S2C). Moreover, if a strong preference or avoidance of AgRP stimulation or a strong correlation of this preference with EPM score were only present at a single phase of estrous, then this would be evident when estimating the maximum or the minimum AgRP preference across the 5 days of testing, but not both. However, both the maximum and minimum preference across the 5 days of testing were strongly correlated with EPM scores in females (Figures S3D-E). These data argue against a dominant contribution of the estrous cycle phase to our findings.

Given that EPM score and sex were predictive of preference for AgRP stimulation after chronic stress, we wondered whether other parameters might provide additional predictive power. We considered the following parameters: the amount of food eaten during AgRP stimulation in the home cage prior to other experiments (Figures S2E-F), body weight measured prior to this initial feeding assay, EPM score before chronic stress (Figures 3C and S3A-C), change in the time spent in the center of the open field chamber before vs. after chronic stress (Figures S2H-I), and preference for AgRP stimulation before chronic stress. Because some of these parameters could be partially correlated, we considered them jointly using a multiple linear regression model. On the last day of chronic stress, 76% of the variance in AgRP preference across females could be explained by this linear model (Figure 3E), while only 36% could be explained across males (Figure 3D). Moreover, the variance explained by this model was significantly higher than for shuffle controls in females but not in males (Figure 3G; black arrows). We confirmed that our model was not overfitting the data by cross-validating with leave-one-out analysis. For both males and females, models for which we omitted a single mouse produced similar fits to the full model that included all mice (Figures 3D-E and S3F-G).

To assess the relative contribution of each parameter, we fit separate linear models for all possible subsets of the five parameters (Figure 3F-G). Strikingly, in female mice, models of AgRP preference that contained parameters related to stress responsivity ranked highest in terms of total variance explained (Figure 3G; right). Specifically, while models including EPM score explained more variance than models including the decrease in time spent in the center zone of the OFT chamber following chronic stress, models including both parameters explained by far the most variance (Figure 3G; right, gray arrows), consistent with the fact that these two parameters were not significantly correlated across females (r = 0.12, *P* = 0.65) (Figure S3H). Similar results were observed on all other days following chronic stress (not shown). Therefore, our results reveal a predictive relationship between the AgRP preferences of individual female mice and several measures of predisposition to stress measured outside of the RTPP paradigm. In contrast, no such relationship was observed in males (Figures 3F and 3H, right). These findings indicate that trait-related sensitivity to stress may explain differences in preference for or aversion to a starvation-like state during chronic stress in female mice.

Additionally, we used a similar multiple linear regression analysis to examine factors predicting inter-individual differences in preference for AgRP stimulation prior to chronic stress. Parameters used for this analysis, all measured before *Phase I*, included the amount of food eaten during the 1-hour AgRP stimulation, the body weight during this session, the EPM score, and the baseline preference for each visual pattern in the virtual environment. In females, such models did not exhibit substantial predictive power (Figures 3E, 3G and 3I; 35% of variance explained, no difference from shuffle). In contrast, the models performed better in males (Figures 3D, 3F, and 3H; 62% of variance explained), largely driven by the relationship between AgRP preference and the amount of food consumed during home cage AgRP stimulation (Figures 3D and 3F; as expected from Figure S2E). This analysis suggests that the efficacy of AgRP stimulation was the dominant factor determining the pre-stress aversiveness of AgRP stimulation in males but not in females. Overall, these analyses support the notion that sex, exposure to chronic stress, and multiple facets of an individual’s predisposition to stress-induced anxiety-like behavior together contribute to how individual mice decide whether to seek a starvation-like state.

### Facial expressions correlate with AgRP stimulation preference after chronic stress

The above findings suggest that, following chronic stress, dwelling in a virtual corridor paired with stimulation of AgRP neurons may be anxiolytic to some female mice, yet aversive or neutral to others. Specifically, anxiety-related traits predicted preferences for AgRP stimulation following chronic stress in female mice. However, such estimates involve averaging behavioral observations across entire sessions. Thus, we considered whether a mouse’s *instantaneous* anxiety-related internal state may also differ when dwelling in a corridor paired with or without AgRP stimulation. We hypothesized that differences in internal state might be reflected in facial expressions, given recent studies that revealed distinct mouse facial expressions during emotionally salient experiences that differ in valence^59–63^. To date, detection of moment-to-moment changes in facial expressions has not been possible in real-time place-preference studies due to the lack of stable, high-resolution facial tracking in freely moving mice. Thus, our head-fixed RTPP paradigm afforded us the unique opportunity to test how facial expressions reflect internal states associated with preference or avoidance of AgRP stimulation. We developed a model-free machine learning-based approach to analyze facial expressions and applied our method to analyze the rich dataset of facial videography collected during each session of head-fixed RTPP behavior in VR (1250 hours across 768 sessions).

We hypothesized that distinct facial expressions might occur during certain instances of dwelling in the presence or absence of AgRP stimulation and that these expressions may be more evident following chronic stress, reflecting a shift in the valence of AgRP stimulation. To test if facial expressions correlated with the state induced by AgRP stimulation, we set out to classify whether, at any given moment, a mouse was experiencing AgRP stimulation, based solely on its facial expression. Thus, to avoid assumptions and enhance sensitivity, we built a classifier using face-related pixels in an image, in contrast to previous studies using human-selected features (e.g., shape of the nose, mouth, or eyes)^63^. Facial contours were estimated using DeepLabCut^64^ and pixels unrelated to the face were excluded (Figure 4A). We built a convolutional neural network (CNN) for each mouse by fine-tuning ResNet-16 (a neural network previously trained on large image datasets)^65^ in order to classify individual frames from either AgRP stimulation or neutral corridors (Figure 4B). We compared the fifth session of RTPP before chronic stress, the fifth session of RTPP following chronic stress, and the two control baseline control sessions prior to AgRP stimulation. We used a balanced number of frames from both stimulation and neutral corridors to avoid biases in classifier training (Figure S4B; Methods). Importantly, we focused on frames in which the mouse was stationary to exclude the confounding influence of locomotion and because our earlier analyses did not reveal strong preferences for either corridor during active exploration (not shown).

**Figure 4.**
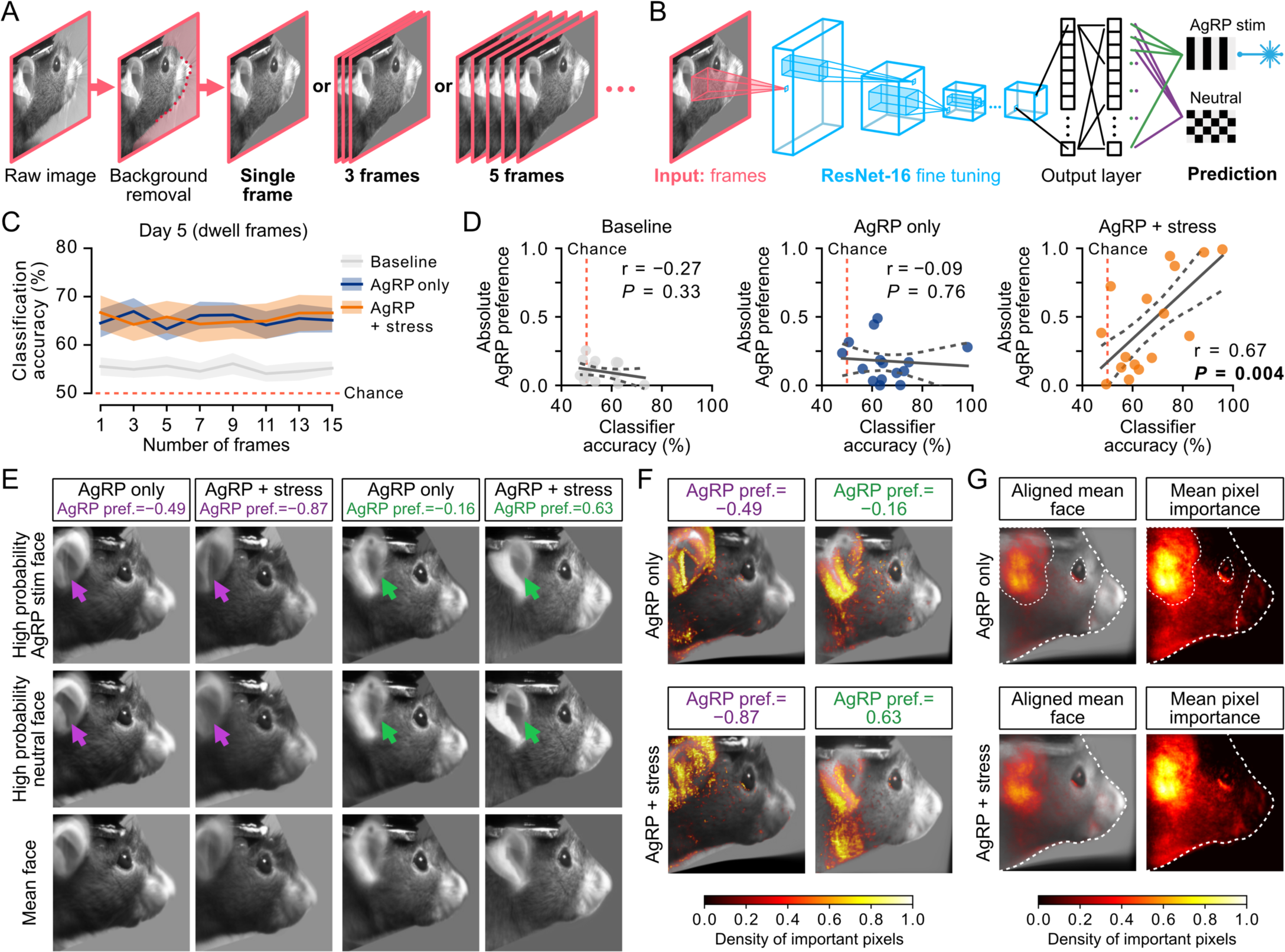
The relationship between AgRP preference and facial expressions. **(A)** Raw images from facial videography were first pre-processed to remove background signals. The area between the contours around the faces, detected using DeepLabCut (see Methods for more details), was removed to reduce the influence of background during classification. Single frames and video clips of increasing size (3-frame and 5-frame clips are shown; 3-15 frames span 110-710 ms; acquired at 20 Hz with 10 ms exposure) were used as input to the neural network for training. **(B)** Schematic of the network trained to predict the corridor associated with each individual facial video image. Images of female mouse faces recorded during the real-time place preference task were first processed to remove background pixels (see also Figure S4SA), and the resulting images (or short movie clips of consecutive images) served as inputs for refining a deep learning model, ResNet-16. In turn, ResNet-16 classifies the input images as those in which the mouse was in the AgRP stimulation corridor or the neutral corridor. Separate networks were trained for each female mouse on Day 5 sessions either prior to or following chronic stress, using matched numbers of frames from both stimulation and neutral corridors. As a control, we analyzed sessions during baseline assessment of cue biases, prior to any AgRP stimulation. **(C)** Mean classification accuracy of each network type based on the number of input frames in a video clip whose duration ranged from a single frame to 15 consecutive frames. All networks from the AgRP stimulation-only condition (blue) and the condition involving AgRP stimulation following chronic stress (orange) performed better than chance (dashed red) and better than the networks trained using baseline control sessions (gray). Data are mean ± SEM. **(D)** The relationship between the classification accuracy for each mouse and the magnitude of behavioral preference for AgRP stimulation during baseline days (gray), and on sessions before (blue) and after chronic stress (orange). r denotes Pearson’s correlation coefficient. Dashed lines represent the 95% confidence band for the linear fit. *P* < 0.05 denoted in bold. (*Baseline*: n=16; *AgRP only*: n=15; *AgRP* + *stress*: n=16 females). **(E)** Comparison of the means of images classified with high certainty (probability ≥ 90%) as occurring in stimulation or neutral corridors. Results are shown for two example female mice, one with aversion (left; purple left) and the other with preference (right; green) for AgRP stimulation, both before and following chronic stress. Purple and green arrows highlight differences in ear position. The number of frames used to calculate the mean images was matched across all conditions. The mean face (bottom row) was constructed using the same number of frames but selected randomly from either corridor. **(F)** Pixel importance maps show the location and density of the pixels that were influential in distinguishing the stimulation faces from the neutral ones (see Methods). Pixel densities are overlaid on aligned mean faces of the same two example mice from (D). The most important pixels aggregate around the ear and eye. **(G)** Mean pixel importance density maps across all female mice before (top row, n = 15 females) and after (bottom row, n = 16 females) chronic stress. Mean maps, overlaid over the aligned mean face from all mice (left) or without the mean face and with higher contrast (right), show the location and density of important pixels. Contours of the face, ear, eye and nose are marked with white dashed lines. See also **Figure S4.**

Using this facial expression analysis, we could determine whether the mouse was in the corridor paired with AgRP stimulation or not during any given frame with an accuracy of 65-70%, which was well above chance (50%). This classification accuracy was near chance levels during the baseline sessions lacking AgRP stimulation, thus ruling out factors unrelated to stimulation (Figure 4C). Consistent with the absence of any noticeable light leakage due to optogenetic illumination (which could interfere with image classification), accuracies in classifying stimulation and no-stimulation frames were near chance levels for frames involving locomotion prior to chronic stress (Figures S4A).

We hypothesized that short video clips might capture the temporal dynamics of facial expressions, and thereby further improve classification. We, therefore, asked whether classifiers trained on 3-frame to 15-frame video clips (110-710 ms, in the range of previously reported mouse behavioral dynamics^66^) improve classification accuracy (Figure 4A-B). However, this was not the case (Figure 4C). Overall, these results indicate that, during some stationary moments but not others, static snapshots of facial expressions can predict whether or not a mouse is dwelling in a location that is paired with a starvation-like state induced by AgRP neuron stimulation.

Having validated our classifier, we asked how each mouse’s facial expressions are influenced by AgRP stimulation after versus before chronic stress. Facial expressions could simply reflect AgRP stimulation-driven reflexive changes in facial expressions, regardless of the stress context or the animals’ preferences for AgRP stimulation. Alternatively, facial expressions could reflect an internal state that is the result of the *interaction* between an animal’s particular reaction to chronic stress and its subsequent change in state upon AgRP stimulation (e.g., relief from anxiety). Consistent with this latter hypothesis, we observed a strong correlation between classification accuracy and the absolute magnitude of AgRP preference, but only after chronic stress (Figure 4D), female mice that strongly preferred or strongly avoided AgRP neuron stimulation both had higher classification accuracy than mice with weaker AgRP preferences (Figure S4C). These findings show that after chronic stress, when more pronounced preferences emerge for corridors associated with or without AgRP stimulation (Figure 2), female mice also begin to display intermittent facial expressions that directly reflect their behavioral preferences for or against a starvation-like state.

To better understand which facial features are important for predicting preference or avoidance of AgRP stimulation, we examined the facial images in greater detail. We first asked how the facial expressions from during dwell frames in the stimulation or neutral corridor differed from the mean facial expression across the entire experiment. Following chronic stress, we observed prominent differences in ear position in a mouse with strong aversion to AgRP stimulation following chronic stress (Figure 4E; right columns), and in a second mouse exhibiting a strong preference for AgRP stimulation (Figure 4E; left columns). The ear positions of both mice differed across corridors, with the ears positioned further back in the absence versus in the presence of AgRP stimulation (Figure 4E; purple and green arrows). This observation motivated us to quantitatively determine which parts of the face contained information related to the differences in facial expressions following chronic stress.

In order to interpret the nature of the information guiding our classifier, we calculated Shapley values for each pixel in each frame using SHAP (SHapley Additive exPlanations) analysis.^67^ This analysis is a common approach for estimating the contribution of a given pixel in guiding image classification (Methods). We calculated the density of important pixels for each mouse for each frame by taking only the top 5% of contributing pixels per frame (Figures 4F-G and S4D). Similar to our observation from the mean images, this analysis revealed that important pixels for an accurate classification were located mainly around the ear and eye for the two example mice described above (Figure 4E). Indeed, in almost all female mice, ear position stood out as an important feature indicating the presence or absence of AgRP stimulation following chronic stress (Figures 4F and S4D). Other features of the face were also informative, though in a less consistent manner across mice (Figure S4D). Overall, these results highlight how rich videography data in head-fixed mice can provide unique insights into behavioral and emotional dynamics related to the interaction of stress and AgRP stimulation in this model of anorexia nervosa.

## Discussion

Individuals who suffer from anorexia nervosa avoid places associated with food and practice ritualized measurements of small amounts of food for consumption, leading to long-term starvation.^3, 4, 8^ Many physiological aspects of anorexia nervosa may be captured by existing mouse models (e.g., ABA).^12, 14^ However, no existing animal study has directly modeled the engagement in behaviors that maintain a starvation state – the most life-threatening of maladaptive behaviors in anorexia nervosa. Here, we provide a new model for understanding the transition to intentional starvation that can often follow from exposure to chronic stress. We find that a subset of female (but not male) mice develops a strong preference for AgRP stimulation after, but not before, exposure to chronic stress. Moreover, the likelihood that a female will develop a preference for this starvation-like state can be predicted by behavioral traits assessed before exposure to chronic stress (e.g., EPM, a measure of anxiety-like behavior).

The emergence of a consistent behavioral preference or avoidance of AgRP stimulation across sessions during chronic stress was mirrored by changes in the way AgRP stimulation influenced facial expressions. Activation of AgRP neurons may defend against starvation in part by decreasing sensitivity to pain/anxiety^29–31, 40^ and thereby promoting foraging. In this way, activation of AgRP neurons via self-starvation may be used as a form of self-medication by certain individuals to mitigate the long-term elevation in anxiety associated with chronic stress. Below, we discuss this new model for studying the willful seeking of AgRP stimulation in the context of previous studies. We then discuss the challenges and benefits of our approach in head-fixed mice for examining this and other understudied psychiatric disorders at the level of facial expressions and high-density imaging and electrophysiology.

### Predisposition to stress may lead to seeking of a star-vation-like

Our findings revealed a pronounced polarization of individual preferences for AgRP stimulation in female mice following chronic stress. In contrast, we did not observe similar stress-driven changes in male mice. This finding points to a clear biological difference between males and females regarding the interaction of stress and hunger. The observation that only a subset of female mice develops a strong preference for AgRP stimulation is akin to the human condition, where not every individual with a stressful life history or trauma develops anorexia nervosa. Furthermore, our findings mirror the 5-10 fold increase in penetrance of anorexia nervosa amongst females in the human population.^6^

Previous studies in humans have suggested that stress-induced changes in food intake and emotional eating can differ between males and females.^68–70^ The type of stress, life history and comorbid conditions also influence the complex interaction between stress and eating. Future studies can exploit the sex-dependent behavioral variability we observed among female mice to investigate the neuronal underpinning. For example, comparing neural activity in the brains of mice that develop a strong preference or aversion for AgRP stimulation after chronic stress is daunting given the large inter-individual differences we observed. However, preselecting female mice with high or low EPM scores prior to running them through our protocol would be an effective strategy for increasing the yield of mice with a strong preference for or aversion to AgRP stimulation. Similar behavioral stratification approaches were recently used to study the dopaminergic system in responses to chronic stress,^71^ and to examine how the mouse insular cortex processes interoceptive signals to maintain balanced levels of fear.^72^ Indeed, this latter study showed that impairment of insular cortex leads to polarization of freezing responses to aversive cues during extinction. Particularly given that insular cortex activity is disrupted in anorexia nervosa,^73–75^ it is tempting to speculate that an impairment in the function of insular cortex following chronic stress may contribute to the polarization of AgRP preferences that we observed.^76^

In the ABA rodent model of anorexia nervosa, animals are food restricted and given access to a voluntary running wheel, leading to a decrease in food intake and an increase in physical activity. These behaviors mimic those observed in anorexia nervosa patients. A recent study^77^ highlighted AgRP neurons as a potential neural target for addressing maladaptive behaviors in anorexia nervosa. Activation of AgRP neurons can mitigate the development and progression of ABA by reprioritizing food intake over stress-induced hyperactivity during periods of food availability.^77^ These findings are consistent with our results that seeking out AgRP stimulation may help displace unpleasant sequelae of chronic stress.

### Possible circuit mechanisms

Like fasting, activation of AgRP neurons leads to a rapid increase in food consumption^24, 25^ and foraging behavior.^29, 30^ However, previous findings suggest that the set of behaviors driven by AgRP stimulation depends on the context. For example, while the activation of AgRP neurons was shown to be mildly aversive in a place preference chamber without food,^26^ AgRP activation is positively reinforcing in a context in which food can be obtained by pressing a lever.^47^ Other studies demonstrated that fasting and stimulation of AgRP neurons are both anxiolytic^29, 30, 40^, drive stereotyped displacement behaviors^29^ in the absence of food, and cause mice to spend more time in the anxiogenic portion of the EPM and OFT chambers (open arm and center zone, respectively).^29, 30, 40^ Similarly, in our task, AgRP stimulation was aversive in males in the absence of food. However, AgRP stimulation was not initially aversive on average in female mice, and some female mice began to strongly prefer stimulation of AgRP neurons when the context became unpredictable and stressful. Thus, our results suggest that in a subset of mice, exposure to chronic stress can change the valence associated with food restriction.

How might AgRP stimulation become reinforcing during chronic stress? Recent studies have shown that chronic stress induces depression-like behaviors by suppressing AgRP neurons, and that chemogenetic activation of AgRP neurons reverses the anhedonia induced by such stress.^78^ Similarly, satiety-promoting arcuate hypothalamic POMC neurons, which become hyperactive following chronic stress, have also been proposed to drive anhedonia.^79^ Thus, AgRP stimulation may counter the effects of chronic stress-evoked hyperactivity of POMC neurons^80^ via direct inhibition of POMC cell bodies and via competition with actions of POMC axons at common downstream targets.

### Facial expressions as a reflection of the effect of AgRP stimulation on internal states

Our analysis of facial videography in head-fixed mice allowed us to detect noticeable yet subtle changes in their facial expressions that may reflect their ongoing emotional state.^59^ Through the application of convolutional neural networks, we successfully classified facial features that were associated with moments of AgRP stimulation in stationary mice. Interestingly, short video clips did not improve classification over static images. As such, additional information may not be contained in the across-frame dynamics. Alternatively, training networks using videos typically requires larger datasets, making them susceptible to overfitting when the available data is limited. In any case, our findings indicate that individual facial snapshots contain enough information to distinguish between different putative internal states.

We found that, after chronic stress, classification accuracy scaled with the amplitude of a mouse’s preference for AgRP stimulation. Internal states, such as relief from anxiety, can result in subtle shifts in facial expressions^59^ that may differ from those associated with acute stressors.^81, 82^ Our facial analyses suggest that chronic stress influences the way facial expressions correlate with the intentional seeking or avoidance of a starvation-like state, even in the absence of immediate stressors. We speculate that these AgRP stimulation-induced changes in facial expressions may reflect inter-individual variability in how intentional seeking of self-starvation may impact moment-to-moment feelings of relief from anxiety after chronic stress. Future research can link these moment-to-moment changes in facial expressions in our task with ongoing neural activity in brain regions that track physiological and interoceptive states, such as the insular cortex,^59, 83^ a region that processes negative emotions in humans^84, 85^ and displays altered activity in patients with anorexia nervosa^73, 74^.

### Future directions

We have developed a head-fixed paradigm that is amenable to conditioned place preference (CPP) and real-time place preference (RTPP) tasks and thus could help elucidate the neural circuits responsible for behavioral preferences across a range of applications. Our head-fixed behavioral paradigm can be coupled with two-photon imaging of single neurons, dendrites and axons, while accurately tracking physiological parameters such as breathing and heart rate. In particular, our approach provides a foundation for future work identifying the neural circuits that underlie the voluntary maintenance of long-term starvation states in individuals with anorexia nervosa.

## Acknowledgements

We thank Brad Lowell, Amelia Douglass, Rachel Essner, Kathryn Evans, Stephen Zhang and Jonna Singh Alvarado for their valuable input on the manuscript, and members of the Andermann lab for useful discussions. We thank Brad Lowell, Amelia Douglass, Rachel Essner, and Yoav Livneh for their critical input throughout the project. We thank Noah Pettit and Christopher Harvey for kindly sharing their design for the virtual reality setup. We thank Kayla Fernando, Praveena Prasad, and Jesseba Fernando for technical assistance and mouse colony maintenance. We thank Dhriti Aiylam for help with data collection. This work was supported by a BBRF Young Investigator Grant (H.K.), a Davis Family Foundation Fellowship, NIH F32 DK112589 and NIH T32 NS007484 (A.L.), R01 DK109930, DP1 AT010971, R01 MH12343, a McKnight Scholar Award, and grants from the Boston Nutrition and Obesity Research Center (P30 DK046200) and the Klarman Family Foundation (M.L.A.).

## Author contributions

H.K. and M.L.A. conceived the study and designed the experiments. A.L. assisted with the initial experimental paradigm design. H.K. and T.P. performed the surgeries. H.K., T.P., M. L., L.P and A.H. performed the experiments. H.K. and O.A. performed the facial videography analysis. H.K. and M.L.A. analyzed the data. H.K. and M.L.A. wrote the manuscript with input from all authors.

## Materials and Methods

### Experimental model and subjects

All mouse care and experimental procedures were approved by the Institutional Animal Care and Use Committee at Beth Israel Deaconess Medical Center. Mice were housed in a 12-hour:12-hour light:dark cycle environment with standard mouse chow and water provided ad libitum. After the animals were returned to their home cage upon conclusion of each experimental session, food was withheld until 30 minutes after the end of the session to prevent the long-lasting effects of AgRP neuron stimulation on food intake.^47^ Mice were singly housed following fiber implantation surgery. Adult male and female mice between the ages of 7–18 weeks were used for this study. *Agrp*-IRES-Cre mice (JAX stock 012899) were crossed with Ai32:RCL-ChR2(H134R)/EYFP (JAX stock 012569) to generate double transgenic mice. No statistical methods were used to determine sample sizes.

### Stereotaxic surgeries

As described in detail below, an optic fiber with a metal ferrule (400 μm diameter core; R-FOC_L400C-39NA; NA 0.39; RWD Life Science) was implanted over AgRP neurons at the midline (AP: −1.6 mm, DV: −5.85 mm, ML: 0 mm) in *Agrp*-IRES-Cre::LSL-ChR2-eYFP mice in order to achieve bilateral stimulation and to maximize activation of AgRP neurons.

Mice were anesthetized with isoflurane mixed in 100% O2 (induction, 3%; maintenance, 1%-2%), and placed into a stereotaxic apparatus (Kopf model 963). After exposing the skull via a small incision, a small hole (∼450-500 μm in diameter) was drilled to allow the passage of the fiber implant. The fiber implant was lowered slowly to the target depth and secured to the skull by cyanoacrylate glue (Loctite) around the ferrule. The fiber implant and a custom-made titanium head-post were fixed to the skull using C&B Metabond (Parkell). For postoperative care, mice were injected intraperitoneally with Meloxicam (0.5 mg/kg). Mice were given at least 14 days to recover before the experiment began.

### Evaluation of the fiber placement

Mice were evaluated for AgRP neuron stimulation-induced food intake at the start of *Phase I* (Figure 1A, gray box) and after the end of *Phase III* (Figure 1A, orange box) in order to ensure the correct placement of the optic fiber.^47^ Mice were habituated in their home cage tethered to an optic fiber patch cord for 30 min one day prior to the experiment. For optogenetic stimulation of AgRP neurons, mice were tethered to an optic fiber patch cord in the home cage and allowed to acclimate for 5 minutes prior to experiments. Mice received photostimulation (473 nm; 10 ms pulses of 20 Hz at ∼11mW; delivered continuously for 1 sec with 3 sec intervals repeatedly^24^) for 1 hour while given access to regular chow. Chow pellets were weighed before and after the photostimulation period. For a subset of mice, food intake during a one-hour period before the photostimulation period was measured (Figure S1C). Experiments were performed during the first 6 hours of the light cycle under ambient light.

Fiber placement was further confirmed by post-mortem histology of the brains. All mice that showed stimulation-induced feeding had the correct placement of the optic fiber over the AgRP neurons expressing ChR2.

A minority of mice (n = 4/36) did not increase their food consumption (<0.3 g) during the hour-long home-cage stimulation protocol (see below) and were thus excluded from subsequent experiments (Figure 1A; *Phase I*).

### Histology and collection of brain samples

Mice were terminally anesthetized with tribromoethanol (Sigma Aldrich) diluted in saline and transcardially perfused with 0.1 M phosphate-buffered saline (PBS, pH 7.4) followed by 10% neutral-buffered formalin (NBF, Fisher Scientific). After removing the fiber implants, brains were extracted. Extracted brains were post-fixed overnight at 4°C in NBF and were cryoprotected in 20% sucrose. Brains were sectioned coronally on a freezing sliding microtome (Leica Biosystems) at 40 μm thickness along the entire length of the arcuate nucleus. Brain sections were mounted on positively-charged slides (Denville Scientific).

Native eYFP fluorescence was amplified with immunohistochemistry as follows. A hydrophobic frame was drawn around the brain sections mounted on the glass slides using a pap pen (Sigma-Aldrich) to contain the solutions during incubation. Slides were washed in 0.1 M PBS and non-specific interactions with the primary antibody were blocked with 3% normal donkey serum in 0.25% TritonX-100 dissolved in PBS for 1 h at room temperature. Sections then were incubated in blocking solution with chicken anti-GFP (1:1,000, Life Technologies, A10262) overnight at 4°C. After slides were washed in PBS 3-4 times for 15 min each round at room temperature, brain sections were incubated in Alexa 488 fluorophore-conjugated secondary antibody (Molecular Probes, 1:1,000) for 2 h at room temperature. Following three rounds of PBS washes, brain sections mounted on the glass slides were covered with Vectashield Antifade Medium, with DAPI (Vector Laboratories). Coverslips (Fisher Scientific) were placed over the sections and sealed with nail polish. Fluorescent images were captured with an Olympus VS120 slide scanner microscope.

### Head-fixed virtual reality setup

Linear virtual reality corridors with alternating cues were designed and rendered in VirRMEn (Virtual Reality Mouse Engine).^86^ Mice were head-fixed on a wheel in front of a screen housed in a laser-cut acrylic box (dimensions; 15 inches in width, 18 inches in height and 21 inches in depth).^87^ The rendered virtual environment was projected by a micro projector (Laser Beam Pro C200) onto a parabolic screen using two reflective mirrors. Mice were able to freely move on the wheel to navigate in the VR environment. Only forward movement was allowed along a linear series of corridors. The forward movement of the mouse was recorded and translated to its position by a custom-made incremental rotary encoder coupled to an Arduino microcontroller. The design of the virtual reality setup and the parabolic screen was based on https://github.com/HarveyLab/mouseVR.

### Virtual reality place-preference

Mice were only exposed to at most one experimental session per day in the virtual reality setup, to minimize any stress due to head-fixation. Each experiment lasted for 35-70 minutes. On the days when the mice received optogenetic stimulation of AgRP neurons, mice were placed in their home cage and food was introduced 30 minutes later to prevent hyperphagia due to delayed effects of AgRP neuron activation on food intake.^47^

Each experiment began with a 5-minute habituation period where mice were allowed to navigate in a corridor with no visual cues. Then, the corridors with the cues were introduced. Two distinct visual cues were presented on the walls of the virtual corridors: horizontal stripes or checkers (Figure 1B). The lengths of the corridors were drawn from an exponential distribution and the total length of either corridor type was matched when averaged across consecutive sets of 10 corridors. Luminance of the visual cues was matched by equalizing the number of light and dark shaded pixels throughout a given corridor. The minimum length of a corridor was empirically determined to ensure that the upcoming cue was visible to the mouse from the previous corridor. A third visual cue, a zig-zag pattern with matching luminance (Figure 1B; orange box), was used for the corridor where the mice received five minutes of tail shocks (Figure 1A; Phase III) in order to prevent aversive conditioning to the cues used in the experiment.

### Conditioning to stimulation/cue pairings

Mice were head-fixed in the virtual reality setup and presented with a corridor with gray walls for 5 min to allow them to habituate. Then, they were teleported into the corridor displaying a single cue. Prior to the beginning of the conditioning experiments involving AgRP stimulation, we assessed each mouse’s bias towards each visual cue presented on the corridor walls. Specifically, for two days, mice could choose between either of the corridors (Figure 1B, gray box). While preferences for either of the cues was close to zero, we did observe interindividual variability in their preferences. Therefore, we accounted for these minor biases by pairing the less preferred cue with the stimulation of AgRP neurons (stim), while the other cue remained unpaired (neutral) (Figure 1B).

Each RTPP session in *Phase II* and *III* was preceded by a series of conditioning sessions where we exposed mice to one of the two cue/stimulus pairings (i.e., the cue paired with stimulation or the neutral cue). On the first day of conditioning, cue stimulus pairing was chosen at random. Mice were exposed to the two cue/stimulation pairings for 6 days in an alternating fashion (3 days per cue/stimulation pairing). During conditioning, mice were allowed to navigate the linear track freely. If the cue on the walls was paired with stimulation (stim cue), blue light (475 nm, ∼11 mW) stimulation was delivered in a similar manner as during the feeding test, for 1 sec every 3 sec at 20 Hz, during the entirety of the time they spent in the corridor.^24, 26^ Stimulation was turned off for sessions where the mouse was in the corridor with the neutral cue. After an initial 10 min of exposure in the corridor with the cue, mice were teleported into a new corridor with gray walls (Figure 1A; *Phase II*) or with the zig-zag pattern where they received tail-shocks (Figure 1A; *Phase III*) for 5 min. This was followed by another 20 min exposure to the cue presented at the start of the session. For the cue paired with stimulation, stimulation was turned off during the initial five-minute habituation period in each session and during the interim five-minute period halfway through the session (Figure 1B; the gray vertical line between the two halves of the session).

### Real-time place preference

Following conditioning to the cue/stimulation pairings, mice were assessed for their preference to or avoidance of AgRP neuron stimulation in head-fixed RTPP. Briefly, mice were allowed to navigate in a linear corridor with alternating visual patterns (Figure 1B). Entry to a corridor with a cue that was paired to AgRP stimulation turned on blue light delivery immediately (473 nm; 10 ms pulses of 20 Hz at ∼11mW; delivered continuously for 1 sec with 3 sec intervals repeatedly). Additionally, we prevented prolonged dwelling in a single cue-associated corridor by teleporting the mice to the next corridor, which contained the opposite cue, if three minutes had elapsed in a single corridor. Depending on which cue the mouse was teleported to, this teleportation triggered optogenetic stimulation (stim cue) or a turning off the stimulation (neutral cue).

We repeated the RTPP for 5 daily sessions with two 30 min blocks per daily session separated by a 5 min break. During this break, mice were either teleported to a gray corridor (serving as sham-stress) or to a corridor with a zig-zag visual cue where they received repeated tail-shock stress. Each day, a single preference index for AgRP stimulation was calculated by summing the total dwell time in each cue corridor across both 30 min blocks.

### Facial videography

Facial videos during the head-fixed behavior were collected using a Flea3 camera (FLIR) equipped with a 50mm fixed focal length lens (Computar, M5018-MP2) and recorded with FlyCapture 2 (FLIR) at a frame rate of 20 Hz. Each frame is triggered by an Arduino board synchronized with VirRMEn^86^ (see Head-fixed virtual reality setup). Exposure of the camera sensor was set at 10 ms. The camera was positioned perpendicular to the mouse’s face. Faces were illuminated with infrared light (850 nm) using an IR lamp (Tendelux).

### Induction of chronic stress

Chronic stress was induced during *Phase III* (Figure 1A; orange box) using repeated mild tail-shocks for 11 days starting on the first day of conditioning and continuing until day 5 of the RTPP experiment. Briefly, mice were teleported into a virtual corridor with a distinct cue (Figure 1B; orange box) where they received mild tail-shocks 20 or 30 minutes after the presentation of the first cue during the conditioning and RTPP days, respectively. Tail-shocks were delivered with two electrode pads (Covidien, Series-S) wrapped around the base of the tail. 50 ms current pulses were delivered at 0.35 mA in bouts lasting for 2-3 secs (12-17 pulses) every 30-35 seconds with a stimulus isolator (Iso-Flex; AMPI). Shock timings were made unpredictable by adding random intervals between each bout and each pulse during a bout.

## Tests to assess anxiety-like behavior

### Elevated Plus Maze Assay

The elevated plus maze (EPM) assay was performed 2-3 days after the first feeding test at the beginning of *Phase I* (Figure 1A; gray box). The EPM was composed of open (exposed to the surroundings) and closed (surrounded by walls of 15 cm height) arms perpendicular to each other (30 cm in length and 6 cm in width), and a center zone (6 cm by 6 cm). The EPM was elevated to 50 cm above the ground. The first 10 minutes after the mouse was introduced to the chamber were used for analysis.

### Open field test

The open field test (OFT) was performed on two days: once before and once after the 11 sessions involving chronic stress (Figure 1A; orange box). Mice were tested in a 40 cm by 40 cm by 40 cm plastic box with an evenly distributed light intensity. They were placed in the center of the OFT chamber. Only the first 10 minutes after the mouse was introduced to the chamber were used for analysis.

### Behavioral analysis

High-resolution videos were captured from above the chamber using a Flea3 camera (FLIR) equipped with a 25 mm fixed focal length lens (Computar, M2514-MP2) and recorded with FlyCapture 2 (FLIR) at a frame rate of 30 Hz. Both behavioral chambers were illuminated under the ambient room lights.

Center positions of the mice were detected using DeepLabCut and used to calculate the time spent in any portion of the experimental chambers.^64^ In EPM, open-arm occupancy was calculated as the ratio of time spent in the open arms over time spent in the closed arms (the center zone was not included as a part of either arm; Figure S2A). In OFT, center zone was designated as an area that is 50% of the total chamber area centered in the middle (Figure S2H). Center zone occupancy was calculated as the ratio of time spent in the center zone over the time spent outside this zone.

## Data analysis

### Identification of the locomotion states

The locomotion state of the animal at each moment in time was defined using a Hidden Markov Model (HMM). We assumed that there are three hidden states of locomotion (see Figure 1C for an example). (1) *Dwell*: when the mouse stays stationary. (2) *Explore*: when the mouse is running or walking fast to explore the virtual environment. (3) *Transition*: where the mouse briefly increases or decreases its speed to transition between a *dwell* state and an *explore* state or between an *explore* state and a *dwell* state. The transition and emission matrices were randomly initialized for the model training. Ζ scored speed values recorded at 10 Hz for all mice and all sessions were used as the model input. The model was trained using the Baum-Welch algorithm (a.k.a. Expectation-Maximization). The most likely sequence of hidden states (a.k.a. locomotion states) were determined using the Viterbi algorithm. The HMM was built using the Python package hmmlearn (version 0.2.8). The resulting states were labeled as dwell, transition or exploration based on their average speed from low to high.

### Preference index

The place preference index was calculated using the time each mouse spent stationary in the VR environment, as follows:

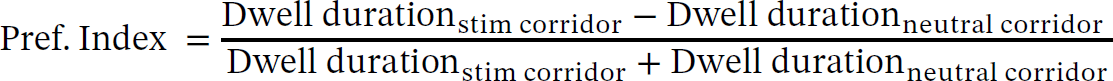

Positive values indicate a positive preference for AgRP stimulation while negative values indicate aversion to AgRP stimulation. A single preference was calculated for each session.

### Correlations between AgRP preference and other variables

A Pearson correlation coefficient (r) was calculated in Python using the SciPy module (version 1.10.0)^88^ in order to test the correlation between AgRP preference and the amount eaten during the feeding test or open arm occupancy in EPM.

Similarly, a correlation matrix was generated by calculating the Pearson correlation coefficient (r) between each possible pair of variables that were used to construct the linear model on Day 5 following stress for female mice (Figure 3).

### Multiple linear regression analysis

The following variables were used in order to determine whether preferences for AgRP stimulation can be predicted on Day 5 following chronic stress: (1) amount of food eaten during 1 hour of AgRP stimulation, (2) weight measured on the day of the feeding test, (3) open arm occupancy score in EPM, (4) change in the center zone occupancy in OFT, (5) baseline preference for the visual cues, and (6) preference for AgRP stimulation prior to chronic stress. We used variables 1, 2, 3, and 5 to predict before-stress preferences and variables 1, 2, 3, 4, and 6 to predict preferences following stress. Separate models were constructed for male and female mice. All variables were normalized prior to constructing the linear models. Ridge regression with L2 regularization was used to estimate the model coefficients. To verify the ability of our models to generalize, we cross-validated the models using leave-one-out analysis. To do so, we constructed individual models after dropping out a single mouse at a time and calculated explained variance (R^2^) using the pooled predictions. All data were preprocessed, and models were constructed in Python (version 3.8) using the scikit-learn package (version 1.2.0).^89^

In order to determine the most predictive variables, we systematically constructed linear models using all possible subsets of the variables mentioned above. Then, we calculated explained variance (R^2^) for each subset. For shuffle controls, the variable labels were shuffled 1000 times for each subset to construct control models and to calculate associated R^2^ values. The contribution of each variable was assessed by calculating the average change in R^2^ values across the subset of models missing that variable as compared to the full model with all variables.

## Classification of facial expressions

### Data preparation and image preprocessing

Frames, when the mouse was stationary (dwell frames) or running (exploration frames), were separated based on the speed of the animal during the facial video recordings (see above). For training the network, we only used dwell frames. Either single static frames or short video snippets of 3 to 15 frames were used to train and analyze the individual networks. These short videos span 110 to 710 ms calculated as follows (frames were recorded at 20 Hz with 10 ms exposure; see Facial videography).

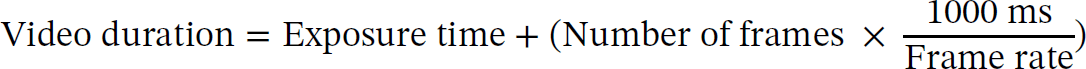

Due to the high correlation between consecutive frames, we split the data into smaller chunks of dwell frames. Thus, there were at least 1000 non-dwell frames between each chunk of dwell frames to minimize the temporal correlation between frames. After dividing the data into chunks, it was split into training and testing data sets, with 80% and 20% of the data allocated to training and testing, respectively. Due to the preference of each mouse for one type of corridor (stimulation or neutral), the number of frames per corridor was unequal. We addressed this bias by matching the total number of stimulation and neutral frames used in both training and testing sets. Specifically, the number of frames from the most preferred corridor was subsampled in order to match the total number of frames from the less preferred corridor.

All input images were preprocessed before training and testing of the neural network. The contours of the face were traced by extracting 14 landmark points surrounding the face (focusing on the nose and the mouth) for each frame using DeepLabCut.^64^ The area that remained outside of the contour points was considered background. The background illumination and any artifacts related to out-of-focus reflectance related to the visual stimuli were removed by replacing the background pixels with a gray value (122 out of a range from 0 to 254 for these 8-bit images). Image pixels were normalized to have a mean of 0.456 and a standard deviation of 0.224.

### Convolution neural network classifier

We built a convolutional neural network (CNN) classifier by finetuning Resnet-16 (a neural network trained with a large image dataset)^90^ in order to identify whether a face came from the corridor paired with AgRP stimulation or from the neutral corridor. Separate networks were trained for each mouse on Day 5 of *Phases II* and *III* (before and after chronic stress) and on Days 1 and 2 of estimation of baseline preference (Figure 1A; *Phase I*). The Resnet-16 network was modified to handle a two-way classification and to accept inputs with more than three dimensions. First, the last layer of the network was replaced with a new linear layer with two distinct outputs, each corresponding to either stimulation or neutral corridors. Second, we expanded the input dimension while multiplying the weights of the original convolutions in order to handle more than 3 frames as input to the network. We used cross-entropy loss with an exponential LR scheduler that decays the learning rate of each parameter group by a factor of 0.75 (gamma) every epoch together with the Adam optimizer^91^ with betas of 0.9 & 0.999. The network training was repeated 20 times (20 epochs) while saving the weights of every epoch. The loss was calculated each epoch and the network with the lowest loss on the valid data was selected. All networks were trained on Nvidia A4000 GPUs. Networks with the best performance were selected for all further analyses for each dataset.

### Evaluation of pixel importance

SHAP (SHapley Additive exPlanations) was used to determine the contribution of each pixel to the classification of frames. The DeepExplainer algorithm from the SHAP package^67^ was used to explain the output of networks we trained for the classification of faces. The entire training dataset was used to train an explainer network for each classifier network. Each frame from the test dataset was then evaluated using this explainer network to obtain Shapley values. A Shapley value estimates the contribution of a pixel to the classification of a given frame as belonging to the AgRP stimulation corridor rather than to the neutral corridor. Therefore, pixels with positive values were considered to be increasing the model’s output (i.e., decision confidence) for the stimulation category, while negative pixels decreased the output. Since our models were constructed for a binary classification (i.e., whether a frame belongs to a stimulation or a neutral corridor), Shapley values of the pixels for either category had identical magnitude but the opposite sign. Thus, we chose to use the Shapley values for the stimulation category only.

In order to determine the important pixels driving classification, we set a threshold for the 95th percentile of the distribution of all Shapley values. Any pixel with an absolute value above this threshold was considered to be important. After determining important pixels for each frame, we calculated a mean value along all the test frames for a given pixel. Resulting density maps were used to identify the spatial facial features that may be important for accurate classification. Before calculating the pixel importance densities, all images were registered to a randomly chosen image frame using the landmark points obtained from DeepLabCut to allow us to align pixels of corresponding facial features.

To align the images of Shapley values so that we could average across frames, for each mouse, we selected a single frame and registered all frames in that session to this frame. We used the landmarks estimated via Deeplabcut to calculate Thin Plate Spline deformation, which we applied to both the mouse facial images and pixel maps of Shapley values.^92, 93^ A similar procedure was used to align pixel maps of Shapley values across mice.

### Statistics

No statistical method was used to predetermine sample size, nor were randomization and blinding methods used. A minority of mice (n = 4/36) that underwent the surgical procedure were excluded from all experiments as they did not increase their food consumption in response to the stimulation protocol (see Evaluation of fiber placement). Two female mice in *AgRP only* and one female mouse in *AgRP + stress* conditions were excluded from the analysis in Figures 4 and S4 due to a technical error during the collection of the facial videos resulting in data loss. Sample sizes (n) for each experiment are reported in the corresponding figure legend.

Statistical analyses were performed using Python (version 3.8) and SciPy module (version 1.10.0)^88^ and are stated in the figure legends. Statistical significance was defined as *P* < 0.05. Whenever applicable, exact *P* values are provided in the figures or figure legends. The data presented fulfill the assumptions of the statistical test employed. A two-tailed paired *t*-test was performed in Figures S1C, S2G, and S2I to determine whether population means are different. A one-tailed *t*-test with Bonferroni correction for multiple comparisons was used in Figure 2E-F only, in order to test whether the AgRP preferences on any of the days prior to stress were less than the baseline preference (predicted based on prior mouse studies and as predicted in our hypothesis based prior human literature; see Introduction) and whether preferences following chronic stress on any of the test days were greater than the preference on the day 5 of prior to stress.

The 95% confidence intervals for individual mice, shown in Figures 2A-2B, are derived through bootstrapping. We separately resampled the dwell durations in the stimulation and neutral corridors 1000 times. The recalculated preference indices across all resampled subsets thus forming the bootstrapped distribution. Null distributions were estimated in Figures S2A-B by resampling the dwell duration in the corridors 1000 regardless of whether they are stimulation or neutral. Similarly, preference indices were recalculated from resampled subsets to construct the null distribution. Preferences that fall outside of the 95% of the null distribution (<2.5^th^ percentile and >97.5^th^ percentile) on a given day were deemed to show significant preference (Figures S2C-S2D). Confidence intervals in Figure 3B were estimated using a leave-one-out method due to the small sample size. Briefly, this involved removing one mouse at a time and subsequently computing the population variance. This was repeated for each mouse and carried out separately for males and for females.

### Data and code availability

The data that support these findings and the code used in the analyses will be made available before the publication of this manuscript. The authors will take the required steps during the revision process to ensure that the data and code are uploaded to a publicly accessible repository.

---

**Figure S1.**
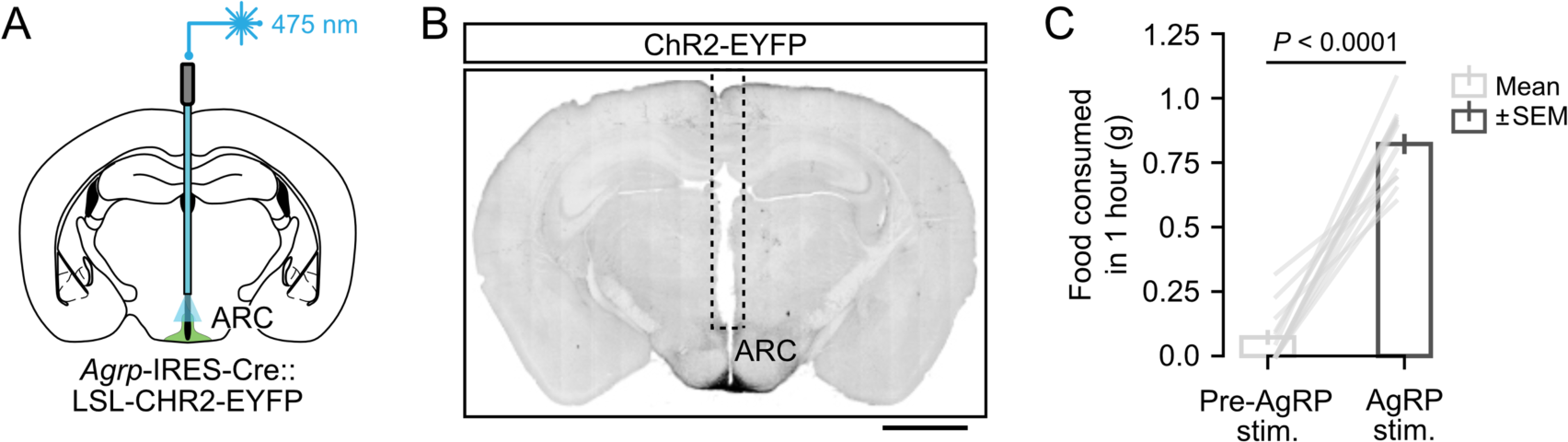
Transgenic targeting of AgRP neurons in the arcuate nucleus of the hypothalamus (related to Figure 1) **(A)** Schematic of optogenetic activation of AgRP neurons in the arcuate nucleus of the hypothalamus (ARC) in an *Agrp*-IRES-Cre::LSL-ChR2-eYFP transgenic mouse line. 475 nm laser light was delivered through a 400 µm optic fiber (see Methods). **(B)** Expression of ChR2-eYFP in AgRP neurons and optical fiber placement above the arcuate nucleus. Scale bar, 1 mm. Image from a representative mouse (n = 15 males, n = 17 females). **(C)** Activation of AgRP neurons leads to increased food consumption during 1 hour of photostimulation. (n = 13 male mice). *P* < 0.0001, two-tailed paired t-test. Data represent mean ± SEM.

**Figure S2.**
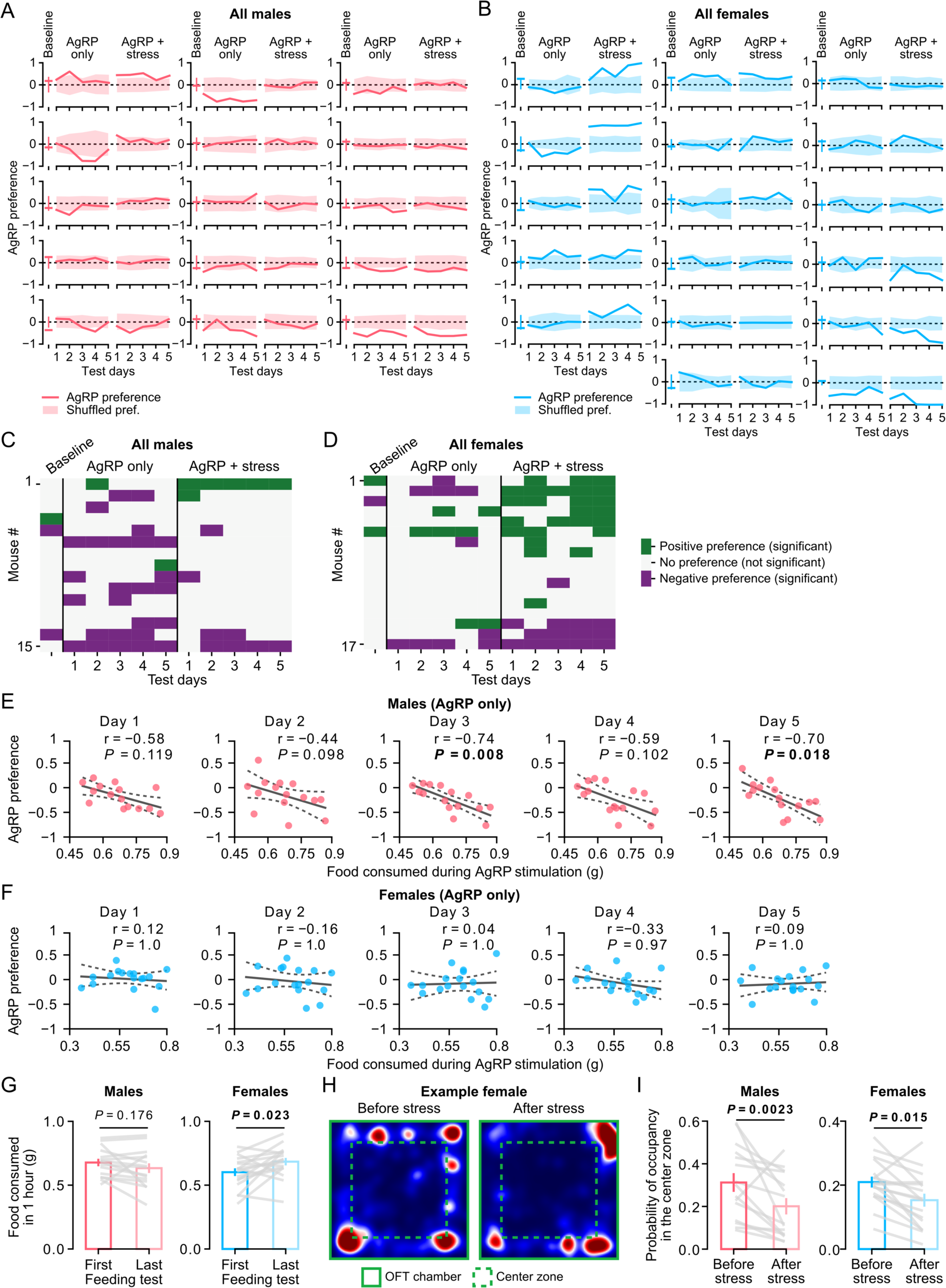
Avoidance of AgRP stimulation in males correlates with the amount of food eaten during the feeding test; Assessment of the efficacy of chronic stress induction (related to Figure 2) **(A-B)** Preference plots for all male (A) and female (B) mice reported in our experiments. AgRP preferences for *Baseline, AgRP only and AgRP + stress* conditions can be compared to the null distributions obtained by calculating a preference index on each day from shuffled dwell durations in either corridor (see Methods for details). All mice are sorted based on their preference on Day 5 of *AgRP + stress*. Shaded areas represent 95% of the range of the null distribution for male (A; n = 15) and female (B; n = 17) mice. **(C-D)** Binarized heat maps of AgRP preference in male (A) and female (B) mice (n = 15 males, n = 17 females), showing the days with significant preferences, defined as those in which the AgRP preference falls outside of the 95% range of the null distributions of shuffled preferences (see panels A-B). **(E-F)** Relationship between AgRP preference and the amount of food consumed during 1 hour of simulation of AgRP neurons on each of the 5 days of RTPP (n = 15 males, n = 17 females) prior to chronic stress (*AgRP only*). Amount of food eaten shows an inverse relationship with AgRP preference in males **(D)**, while no such relationship was observed in females **(E)**. r denotes Pearson’s correlation coefficient; dashed lines represent the 95% confidence band for the linear fit. *P* < 0.05 denoted in bold; *P* values are corrected for multiple comparisons using Bonferroni correction. Each circle represents a single mouse. **(G)** The amount of food eaten in response to 1 hour of stimulation of AgRP neurons in males (left) and females (right) on the first day of Phase I (Figure 1A, middle) and at the end of Phase III (Figure 1A, right) (n = 15 males, n = 17 females). *P* < 0.05 denoted in bold; *P* = 0.176 (males), *P* = 0.023 (females), Data represent mean ± SEM. **(H)** Heat map of occupancy in the open field test (OFT) chamber for an example female mouse before and after chronic stress. Boundaries of the chamber and the center zone (50% of the total area) are denoted by solid and dashed green lines, respectively. **(I)** Occupancy in the center zone for males (left) and females (right). Mice of either sex spent less time in the center zone of the OFT chamber after stress (n = 15 males, n = 17 females). *P* < 0.05 denoted in bold; *P* = 0.0023 (males), *P* = 0.015 (females), two-tailed paired t-test.

**Figure S3.**
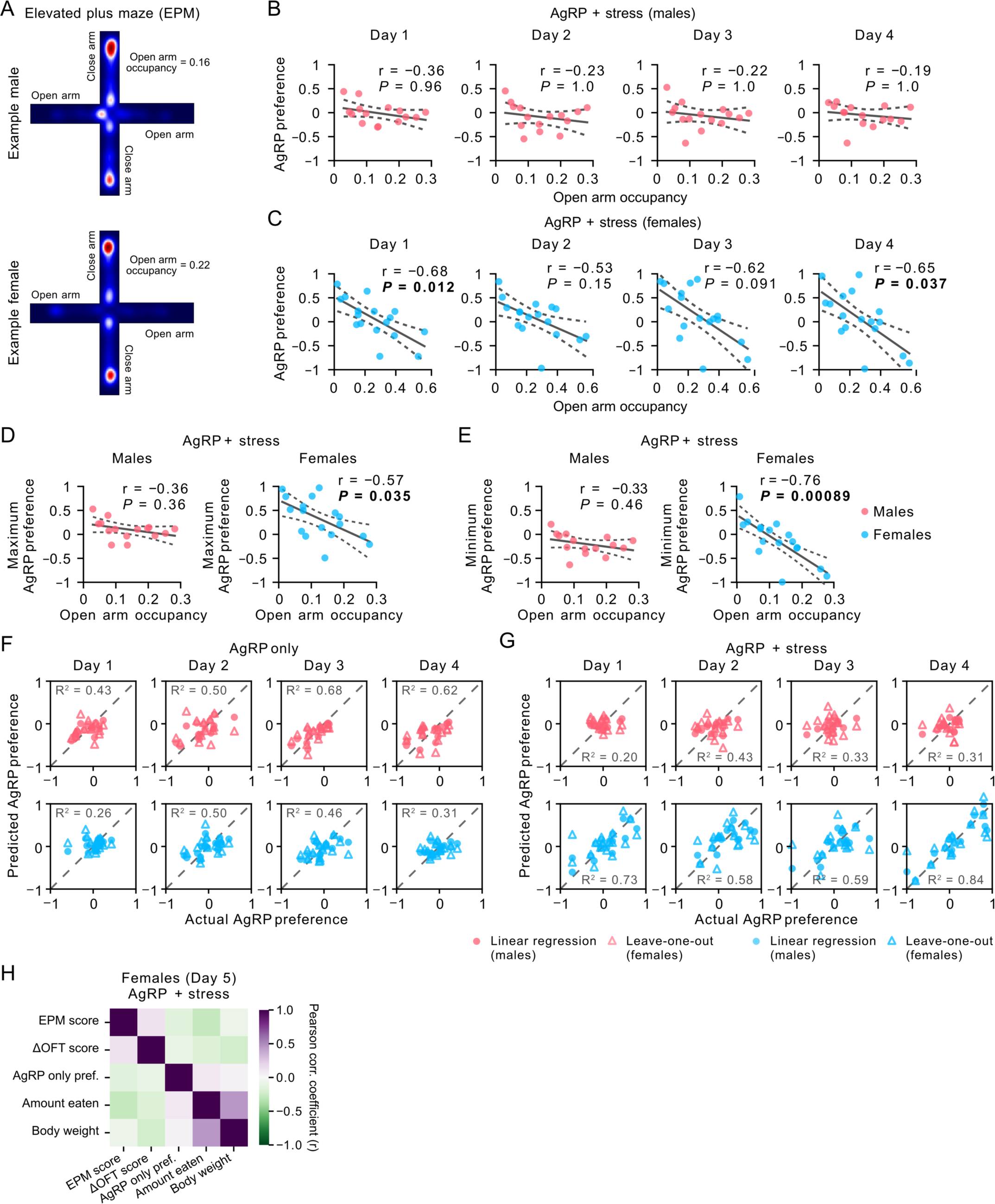
Open arm occupancy in EPM, a measure of predisposition to stress, correlates with preference for AgRP stimulation following stress in females (related to Figure 3) **(A)** Example heat maps of occupancy in the elevated plus maze (EPM) assay from an example male (top, open arm occupancy score = 0.16) and female (bottom, open arm occupancy score = 0.22) mouse. EPM scores were calculated as the ratio of time spent in the open arm (aligned horizontally) over the closed arm (aligned vertically). **(B-C)** The relationship between open arm occupancy and AgRP preference following stress on the first 4 days of the RTPP experiment for male (B) and female (C) mice (n = 15 males, n = 17 females). Each filled circle represents a single mouse. r denotes Pearson’s correlation coefficient. Dashed lines indicate the 95% confidence band for the linear fit. *P* < 0.05 denoted in bold; *P* values are corrected for multiple comparisons using Bonferroni correction. Each circle represents a single mouse. **(D-E)** The relationship between open arm occupancy and maximum (D) or minimum (E) preference across days, shown for female mice following chronic stress (n = 17 females). See also legend of Figures 3D-E. r denotes Pearson’s correlation coefficient; dashed lines represent the 95% confidence band for the linear fit. *P* < 0.05 is indicated in bold. Each circle represents a single mouse. **(F-G)** Scatter plots of actual preference for AgRP stimulation and predicted preference estimated from a multiple linear regression model before chronic stress (*AgRP only*) (F) and after chronic stress (*AgRP + stress*) (G) for male (top) and female (bottom) mice for Days 1 to 4. Each point represents a single mouse estimate from the multiple linear regression (filled circle) and from the leave-one-out analysis (open triangle) (n = 15 males, n = 17 females). **(H)** Cross-correlation matrix of 5 variables used in the full linear model and in linear models with subsets of variables in Figures 3E, 3G, and S3G. r: Pearson’s correlation. None of the variable pairs were significantly correlated (all *P*’s > 0.05).

**Figure S4.**
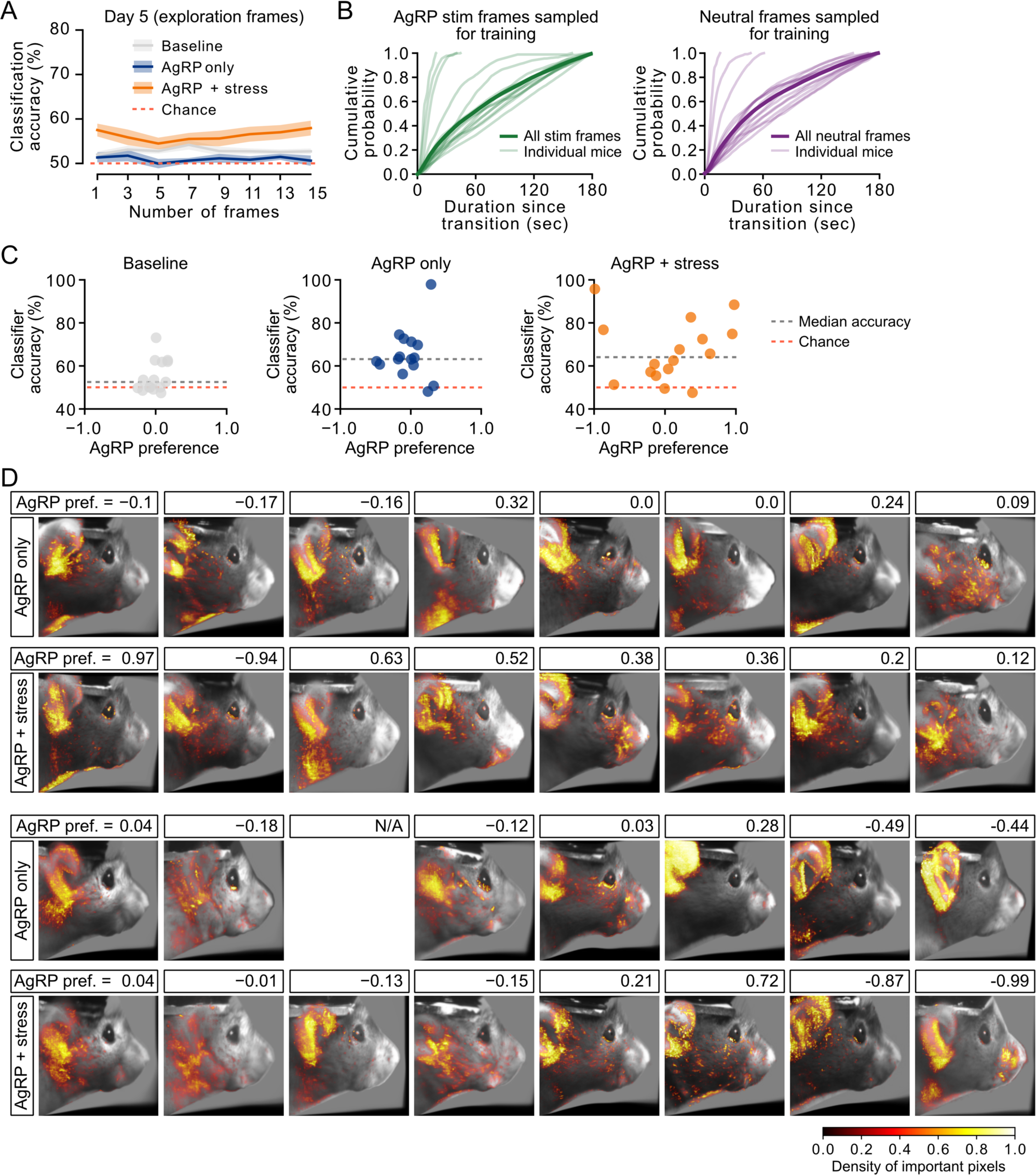
Discriminability of facial expressions is decreased during locomotion; Pixel importance maps of all individual female mice (related to Figure 4) **(A)** Cumulative probability of using a frame that occurred at a given delay after transitioning into a new stim or neutral corridor. Frames used in training the classifiers were randomly sampled for each cue, starting from the entry into a corridor until the mouse exited the corridor. **(B)** Mean classifier accuracy was generally lower on frames in which the mouse was running as compared to frames when the mouse was stationary (compare to Figure 4B), on baseline days (gray), on AgRP stimulation sessions prior to stress (blue), and on AgRP stimulation sessions after stress (orange). Red dashed line: chance performance. Data represent mean ± SEM across female mice. **(C)** The relationship between the preference for AgRP stimulation and the classifier accuracy of each mouse during baseline days (gray), before chronic stress (blue), and after chronic stress (orange). The two example mice shown in Figure 4D-E are selected among mice with AgRP preferences above the median classification accuracy in the AgRP + stress condition. r denotes Pearson’s correlation coefficient; dashed lines represent the 95% confidence band for the linear fit. *P* < 0.05 denoted in bold. **(D)** Pixel importance maps showing the location and density of the pixels that were influential in classification for each female mouse, prior to stress (rows 1 and 3) and following chronic stress (rows 2 and 4). Density maps are overlaid over aligned mean faces during stimulation frames from individual mice. Mice are sorted by their preference for AgRP stimulation on Day 5 following chronic stress.

